# Engineering bacteria for biogenic synthesis of chalcogenide nanomaterials

**DOI:** 10.1101/266502

**Authors:** Prithiviraj Chellamuthu, Frances Tran, Kalinga Pavan T. Silva, Moh El-Naggar, James Q. Boedicker

## Abstract

Microbes naturally build nanoscale structures, including structures assembled from inorganic materials. Here we combine the natural capabilities of microbes with engineered genetic control circuits to demonstrate the ability to control biological synthesis of chalcogenide nanomaterials in a heterologous host. We transferred reductase genes from both *Shewanella* sp. ANA-3 and *Salmonella enterica serovar Typhimurium* into an heterologous host (*Escherichia coli)* and examined the mechanisms that regulate the properties of biogenic nanomaterials. Expression of arsenic reductase genes and thiosulfate reductase genes in *E. coli* resulted in the synthesis of arsenic sulfide nanomaterials. In addition to processing the starting materials via redox enzymes, cellular components also nucleated the formation of arsenic sulfide nanomaterials. The shape of the nanomaterial was influenced by the bacterial culture, with the synthetic *E. coli* strain producing nanospheres and conditioned media or cultures of wild type *Shewanella sp.* producing nanofibers. The diameter of these nanofibers also depended on the biological context of synthesis. These results demonstrate the potential for biogenic synthesis of nanomaterials with controlled properties by combining the natural capabilities of wild microbes with the tools from synthetic biology.

## Introduction

Biological synthesis of nanomaterials provides exciting new routes to synthesize materials for use in environmental remediation, automobiles, photovoltaics, aircrafts, medical imaging, and medical implants (Thakkar *et al.*, 2010; Hulkoti and Taranath, 2014). Biogenic synthesis of nanomaterials is a process which uses biological components from bacteria, fungi, plants, and viruses to synthesize nanomaterials. Microbes found in nature are capable of building nanoscale materials, such as magnetsomes and silicates (Cai *et al.*, 2007; Yan *et al.*, 2012). Microbes have also evolved redox enzymes capable of processing many starting materials used in semiconductor and metallic nanostructure, such as manganese, iron, selenium, arsenate, uranium, and chromium (Nealson *et al.*, 2002; Liu *et al.*, 2002; Yan *et al.*, 2012). These capabilities suggest the possibility of large scale synthesis of nanomaterials by cells, similar to the large scale biogenic synthesis of organic compounds including biofuels and drugs (Cai *et al.*, 2007; Yan *et al.*, 2012). Cellular construction of nanomaterials can also leverage the continued advancement of genetic control circuits, enabling the combination and fine-tuned control of synthesis pathways to modulate nanomaterial properties.

Nanomaterials, such as cadmium sulfide, arsenic sulfide, gold nanoparticles, silver nanoparticles, and zinc oxide have previously been synthesized using biological microorganisms (Narayanan and Sakthivel, 2010; Quester *et al.*, 2013; Jacob *et al.*, 2016; Plaza *et al.*, 2016). These materials include both nanomaterials composed of single elements, such as nanoparticles synthesized from gold or silver, as well as nanomaterials composed of multiple elements, such as cadmium sulfide, arsenic sulfide and zinc sulfide. For instance, arsenic sulfide nanofibers with semiconductor properties were synthesized by wildtype *Shewanella* species (Lee *et al.*, 2007; Jiang *et al.*, 2009; McFarlane *et al.*, 2015). Gold nanoparticles and CdTe/CdS quantum dots have even been patterned using a biological template (Chen *et al.*, 2014). A major challenge in making nanomaterials is the control of nanomaterials properties, including size, shape, and composition. Recent reports show that microbes influenced the size of nanoparticles (Flenniken *et al.*, 2004; Sweeney *et al.*, 2004; Bai *et al.*, 2006; Bakhshi and Hosseini, 2016). Synthetic biology offers new strategies to control the properties of nanomaterials by regulating the expression levels of various biochemical pathways that could control the specific attributes of nanomaterials.

In this work, we take advantage of the ability of microbes to change the oxidation state of metals and metalloids. Microbes have evolved both specific and non-specific enzymes to transfer electrons to metals such as iron and manganese as well as metalloids such as selenium and arsenic. These redox reactions are either part of the respiratory chain or a detoxification pathways. Here, we transfer redox pathways from two strains of bacteria, *Shewanella sp.* ANA-3 and *Salmonella enterica serovar Typhimurium*, into a heterologous host to synthesize arsenic sulfide nanomaterials. Arsenic and thiosulfate reductase genes were cloned into plasmids and expressed under the control of inducible promoters (Bang *et al.*, 2000b). Using this synthetic system, we investigated how the properties of synthesized nanomaterials depend on biological activity, finding that both redox activity and nucleating factors produced by the cells control the size and shape of the resultant nanomaterials.

## Results

### Engineering *E. coli* to reduce arsenate and thiosulfate

To create a synthetic microbial system capable of producing arsenic sulfide nanostructures, genes for arsenic and thiosulfate reduction were introduced via plasmids (Table 1) into an *E. coli* host. For arsenic reduction, *arsDABC* (4.2 kb) and *arrAB* (3.2 kb) from *Shewanella* sp. ANA-3 were assembled into two plasmids. *arrAB* is an arsenic operon giving *Shewanella* sp. ANA-3 the capability to use arsenate As(V) as a terminal electron acceptor. The *arsDABC* operon is a separate arsenic detoxification pathway from the same strain (Saltikov and Newman, 2003; Saltikov *et al.*, 2003; Malasarn *et al.*, 2008). Both pathways reduce arsenate As(V) to arsenite As(III)—necessary for formation of arsenic sulfide nanomaterial. For thiosulfate reduction, thiosulfate reductase gene (*phs* ABC)(Bang *et al.*, 2000b) from *Salmonella enterica serovar Typhimurium* was obtained from Addgene. Previously, researchers used this plasmid in an engineered system for bioremediation applications (Bang *et al.*, 2000a, 2000b). Plasmid maps (Figure S1) and primers used (Table S1) are available in the SI.

**Table 1:**
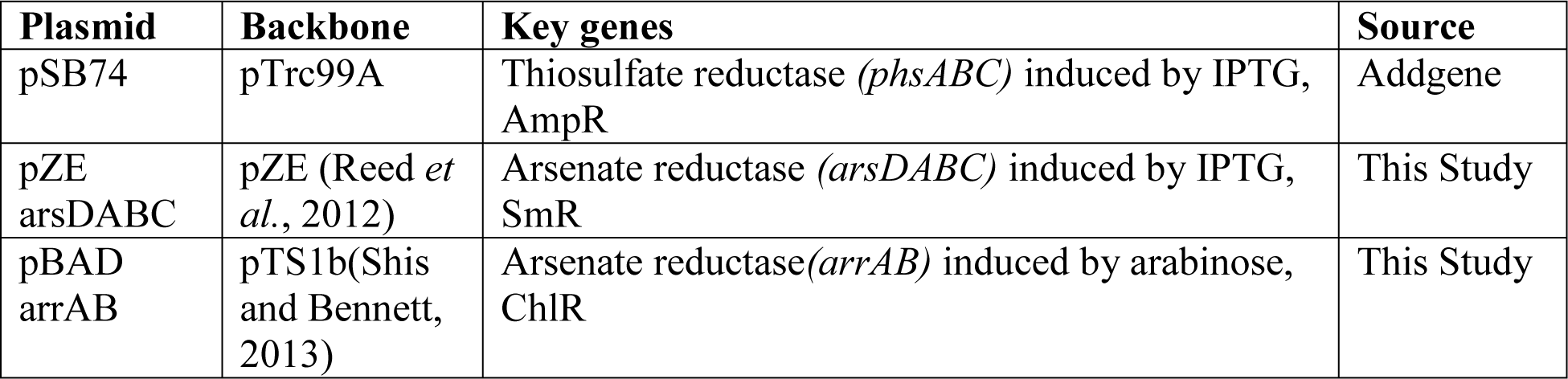
List of plasmids used in this study.

| Plasmid | Backbone | Key genes | Source |
| --- | --- | --- | --- |
| pSB74 | pTrc99A | Thiosulfate reductase (*phsABC*) induced by IPTG, AmpR | Addgene |
| pZE<br>arsDABC | pZE (Reed *et al.*, 2012) | Arsenate reductase (*arsDABC*) induced by IPTG, SmR | This Study |
| pBAD<br>arrAB | pTS1b(Shis and Bennett, 2013) | Arsenate reductase(*arrAB*) induced by arabinose, ChlR | This Study |

Figure 1A shows a schematic of engineered *E. coli* strains involved in thiosulfate reduction to sulfide and arsenate reduction to arsenite, two substrates necessary for arsenic sulfide production. The strains were cocultured for experiments involving the biogenic synthesis of arsenic sulfide nanomaterials.

In control experiments, *E. coli* without the plasmids were tested for the ability to reduce arsenate and thiosulfate. As shown in Figure 1B, after 24 hours, *E. coli* with plasmid pSB74 reduced 2.5 mM thiosulfate to 350 µM sulfide, ten fold more than the no plasmid control. Similarly, arsenate reduction capabilities of *E. coli* strains with plasmids pZE arsDABC and pTSlb arrAB were quantified, as shown in Figure 1C. When induced, strains carrying the arsenic reduction plasmids were able to reduce 5 mM arsenate to 3.8 mM arsenite at 72 hours. Uninduced cultures generated 1.6 mM arsenite after 72 hours. Background levels of arsenate reduction in cells without the plasmids produced 0.8 mM arsenite.

### Synthesizing arsenic sulfide nanomaterials with engineered *E. coli*

We tested the ability of engineered *E.coli* cells with inducible thiosulfate reductase and arsenate reductase to synthesize arsenic sulfide nanomaterials. After induction of the reductase genes, arsenate and thiosulfate were added to the culture. *E. coli* cells expressing reductase genes formed yellow precipitate after 12-14 hours, indicating the formation of arsenic sulfide material. Arsenite reacts with sulfide to form arsenic sulfide (As_2_S_3_) nanomaterial. In control experiments, WT *E. coli* cultures not expressing the reductase genes did not form any precipitate even after 72 hours, as shown in Figure 2A. Although sulfide alone has been shown to reduce As(V) to As(III), in control experiments shown in Figure S3, this low level of arsenic reduction was not sufficient to generate any precipitate.

**Figure 1:**
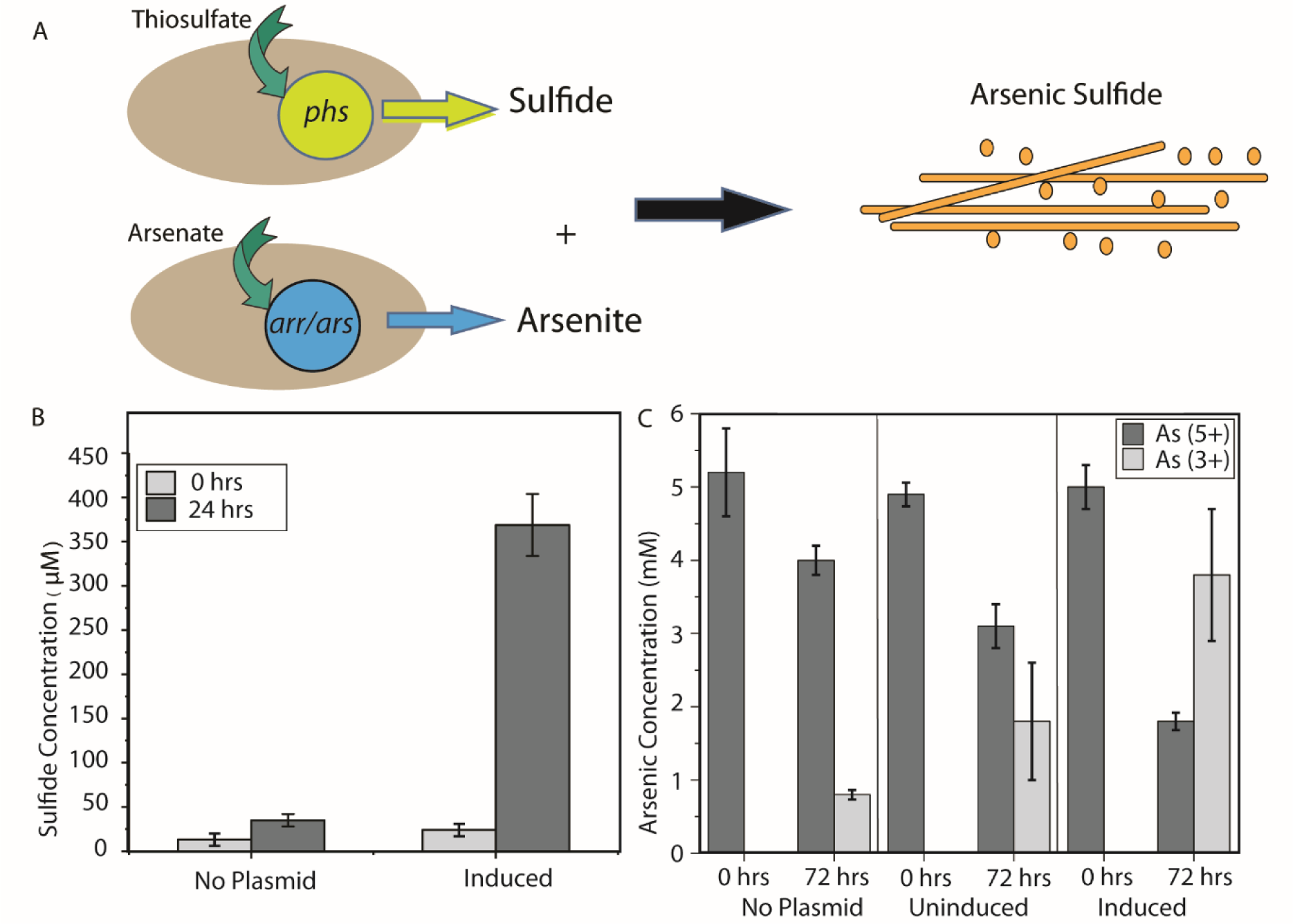
Reduction of thiosulfate and arsenate by engineered *E. coli*. A) Plasmids containing pathways for the reduction of arsenate (*arr/ars*) and thiosulfate (*phs*) were inserted into *E. coli* host cells, enabling the formation of arsenic sulfide nanomaterials. B) *E. coli* expressing the thiosulfate reductase produced high concentrations of sulfide. C) Expression of arsenic reduction enzymes from *Shewanella sp.* in *E. coli* increased the rate of arsenite production as compared to negative controls in uninduced cells or cells without the plasmid.

The precipitate morphology was assessed using scanning electron microscopy. The engineered *E. coli* cultures formed spherical arsenic sulfide structures with an average diameter of 381+/- 246 nm (Figure 2B), which was surprising given that arsenate and thiosulfate reduction by *Shewanella sp.* ANA-3 is known to synthesize nanofibers of arsenic sulfide, as shown in Figure 2C and previously reported (McFarlane *et al.*, 2015). The shape of the material was influenced by the microorganism expressing the reductase genes. Energy dispersive X-ray spectroscopy was used to characterize the elemental composition of the precipitate from both *E. coli* and ANA-3 cultures. The precipitates had signature peaks (Figure S2) corresponding to arsenic and sulfur, confirming the yellow precipitate for both host strains was arsenic sulfide. Table 2 summarizes the nanomaterials formed for the conditions and strains described above.

**Figure 2:**
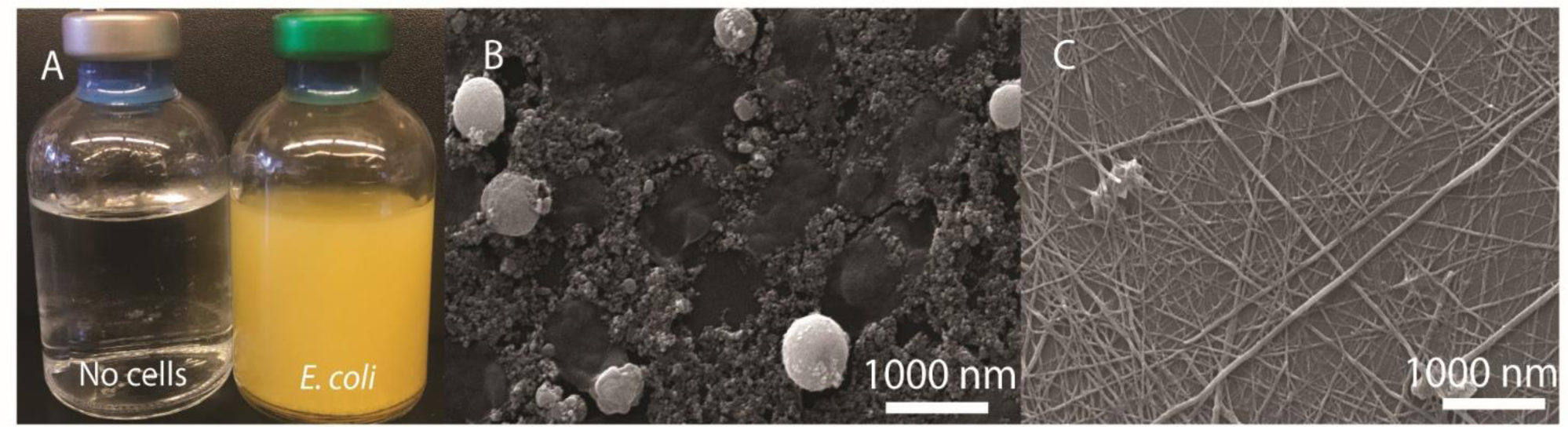
Biogenic production of arsenic sulfide nanomaterials. A) Bottles containing engineered *E. coli* expressing thiosulfate and arensate reductases form yellow precipitate, whereas control bottles without cells remain clear. Scanning electron micrograph images of nanomaterials produced by engineered *E. coli* (B) and wildtype *Shewanella sp.* ANA-3 (C). Both samples were collected at 72 hours.

**Table 2:** Characteristics of biogenic nanomaterials formed through celluar reduction of arsentate and thiosulfate.

| Cell type | Reactants | Precipitation | Nanomaterial | Size (nm) |
| --- | --- | --- | --- | --- |
| *Shewanella* ANA-3 | arsenate, thiosulfate | yes | fiber | 46+/-14 |
| *E. coli* with pZE arsDABC and pBAD arrAB and *E. coli* with pSB74 | arsenate, thiosulfate | yes | spheres | 411+/-125 |

### Cellular components were needed to induce formation of arsenic sulfide nanomaterials

Control experiments were run to determine what roles the cells played in nanomaterial synthesis other than the reduction of the starting arsenate and thiosulfate. To identify additional roles of bacterial cells in nanomaterial formation, reduced substrates, 5 mM arsenite and 10 mM sulfide, were added to cultures of cells and media without cells. Bottles containing minimal media did not form precipitate (Figure 3A and B), indicating a role for cellular components in the nucleation of arsenic sulfide nanomaterial. When arsenite and sulfide were added to anaerobic bottles with *E. coli*, we observed precipitation of arsenic sulfide material within hours (Figure 3D). Cell-free supernatant was also tested, which also initiated nanomaterial synthesis (Figure 3C). Unlike the arsenic sulfide spheres formed when arsenate As(V) and thiosulfate were added to cultures of engineered *E. coli*, the addition of the reduced substrates to *E. coli* cultures formed nanofibers, as shown in Figure 3D.

To further quantify the formation of arsenic sulfide material in these contexts, we measured absorbance at 375 nm, as in previous work (McFarlane *et al.*, 2015). As shown in Figure 3, in abiotic media no precipitate was observed, but absorbance at 375 nm increased as compared to samples with no arsenite or sulfide added. This result suggests that smaller particles of arsenic sulfide form in the absence of cellular components, but do not form a visible precipitate. Bottles containing cells or cell supernatant formed precipitate and had a larger increased in absorbance at 375 nm.

**Figure 3:**
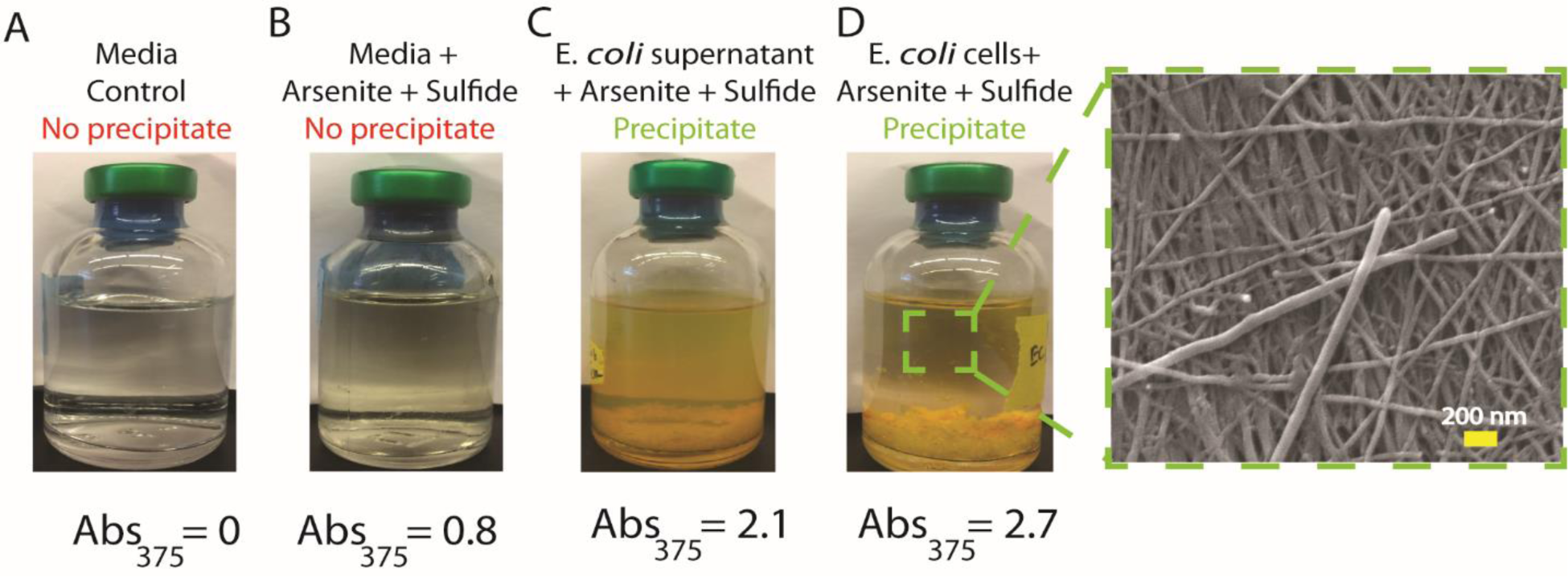
Cellular components were needed to precipitate arsenic sulfide nanomaterials. A) Control bottle of media with no arsenite and sulfide formed no precipitate. B) Bottles with media, arsenite, and sulfide formed no precipitate. Absorbtion of 375 nm light indicates some arsenic sulfide material in solution. C,D) Bottles with arsenite and sulfide containing either cell-free supernatant or *E. coli* cells produced solutions with high absorbance at 375 nm and large amounts of yellow precipitate, characteristic of arsenic sulfide materials. SEM images revealed arsenic sulfide nanofibers were produced when arsenite and sulfide were added to *E. coli* cultures.

We also tested the heat sensitivity of the cellular compounds involved in nanomaterial precipitation. Our results show that the cellular components involved in nucleation were not heat sensitive. Both autoclaved cell culture and autoclaved supernatant were capable of inducing precipitate formation. A summary of results and conditions tested with direct addition of arsenite and sulfide are summarized in Table S3. These results show that biomolecules secreted by the cells are necessary for the initiation of arsenic sulfide material precipitation. Unlike other metals, such as iron and cadmium which readily react with sulfide to form iron sulfide and cadmium sulfide complexes, under our experimental conditions arsenic sulfide requires a heat-stable nucleating agent provided by cell.

Additionally, we tested which conditions were sufficient for nucleation of arsenic sulfide nanomaterials. We then tested if the or ganic molecules of biological origin found in a rich, complete media could induce precipitate formation, specifically LB, which contains yeast extracts, and minimal media with vitamins (see Table S2 for media components). These complex media also did not initiate arsenic sulfide precipitation. Results from abiotic experiments are summarized in Table 3.

**Table 3:** Abiotic conditions tested for arsenic sulfide precipitation.

| Abiotic Conditions | Reactants | Precipitation | Nanomaterial | Size (nm) |
| --- | --- | --- | --- | --- |
| Buffer solution | arsenite, sulfide | no | NA | NA |
| LB media | arsenite, sulfide | no | NA | NA |
| Minimal media with minerals and vitamins | arsenite, sulfide | no | NA | NA |

### Size dependence of arsenic sulfide nanofibers on synthesis conditions

Finally, we assessed whether biotic synthesis conditions influenced nanostructure dimensions. We compared nanofibers synthesized when arsenite, As(III), and sulfide were added to solutions of cells, cell-free supernatant, autoclaved cells, and autoclaved cell-free supernatant for both *E. coli* and *Shewanella sp.* ANA-3. SEM images of nanofibers from each condition were analyzed using ImageJ (Abràmoff *et al.*, 2004) to extract the diameter of each nanofiber.

Solutions derived from *E. coli* cultures tended to form wider nanofibers than solutions derived form *Shewanella sp.* ANA-3 cells. Although fibers from all conditions were similar in width, statistical analysis revealed significant shifts in fiber widths for some conditions (Table S4 and S5). In general, the size variation was larger for fibers produced in *E. coli* conditioned media (Figure 4 and Table S3). These results suggest that biogenic synthetic systems may be able to control nanostructure properties including size.

**Figure 4:**
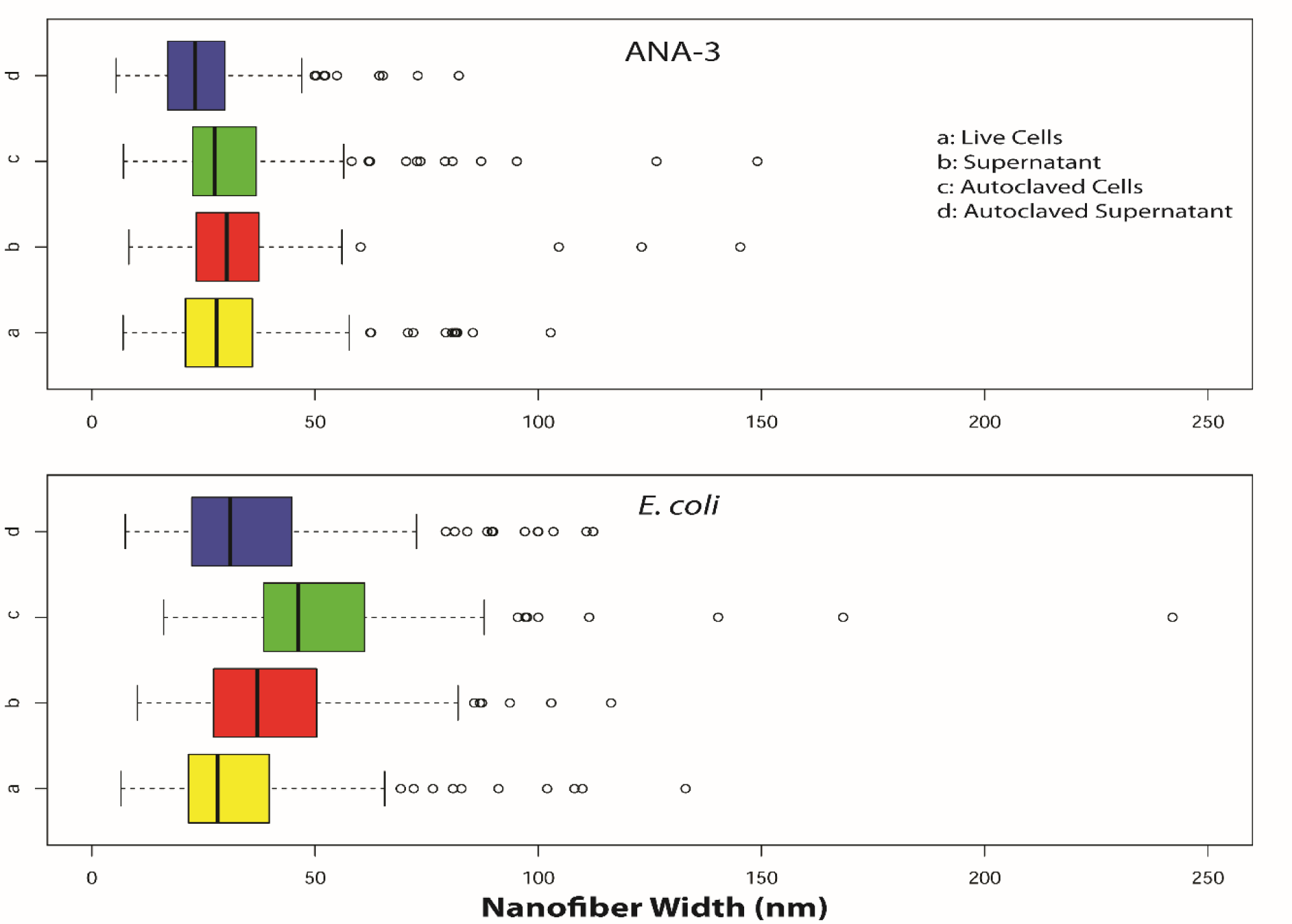
Size distribution of arsenic sulfide nanowires synthesized by the addition of arsenite and sulfide to cellular solutions derived from cultures of WT *Shewanella sp.* ANA-3 (top) or engineered *E. coli* (bottom). Nanomaterials dimensions were calculated from SEM images.

## Discussion

We implemented synthetic gene circuits to synthesize biogenic nanostructures. Reductases from both *Shewanella sp.* ANA-3 and *Salmonella enterica* were expressed in a heterologous *E. coli* host. Expression of these reductases enabled cultures of *E. coli* to process thiosulfate and arsenate and produce sulfide and arsenite—precursors for synthesis of arsenic sulfide nanomaterials. The ability of bacteria to influence the production of nanomaterials has been reported previously(da Costa *et al.*, 2016), and here we explore how several biological parameters influenced the properties of the synthesized nanostructures.

For the formation of arsenic sulfide nanomaterial under ambient conditions, the presence of yet unknown biological components involved in nucleation were critical for material synthesis. In the absence of cells or cellular components, arsenic sulfide nanostructures did not precipitate out of solution. This points to an essential role for biological compounds in nucleating nanoscale materials. Nanomaterial nucleation is an active area of research, as understanding nucleation is critical to control the homogeneity of nanomaterials (Thanh *et al.*, 2014). In some biological systems the nucleating agent has been identified. Cadmium sulfide nanoparticles can nucleate on a cystathionine y-lyase enzyme (Dunleavy *et al.*, 2016), and other known nucleation materials including enzymes, peptides and cells extracts have been reported (Merzlyak and Lee, 2006; Spoerke and Voigt, 2007; Varma, 2012; Seker *et al.*, 2017). For arsenic sulfide nanomaterial, the nucleating agent can be provided by either *E. coli* or ANA-3 cells. We even found the supernatant of these cultures to be sufficient for nucleation of nanomaterial, and that the nucleating agent remained active after autoclaving the solution. The nucleation agent however was somewhat specific to these cultures. Media containing biological components such as digested proteins and vitamins did not precipitate material. These results demonstrate that microbes have multiple roles in nanostructure formation, not only processing starting materials, but also regulating the nucleation of materials.

Besides providing nucleation centers, the biological context of synthesis also influenced both the size and shape of arsenic sulfide nanomaterials. When substrates necessary for arsenic sulfide nanomaterials were processed by engineered *E. coli* spherical nanoparticles were formed, whereas all other conditions produced the nanofibers reported in previous studies (Lee *et al.*, 2007; McFarlane *et al.*, 2015). The switch to spheres was not due to nucleation of the material, as the direct addition of arsenite and sulfide to live *E. coli* cells with or without the reductases resulted in the nanofibers. The mechanism that regulates nanoparticle shape was not determined. Previous work showed the influence of the organism on material shape and size (Li *et al.*, 2011; Dahoumane *et al.*, 2017). Gold nanoparticles made by *Shewanella* were spherical (∼12 nm) in shape (Suresh *et al.*, 2011); however, *E.coli* produced triangles and hexagons (Du *et al.*, 2007). This demonstrates organisms have an interesting role in controlling the shape and size of the nanomaterials. Additionally, the available concentrations of arsenite and sulfide, modulated by the expression level of the reduction genes, could be another potential reason for this variation in shape. Differences in the nanostructure dimensions were also dependent on the biological context of synthesis (Dahoumane *et al.*, 2017). As shown in Figure 3, fibers derived from *E. coli* cultures were slightly wider and more heterogeneous. Statistical comparative analysis (Student T-test) on nanofiber width among these conditions are presented as tables in SI (SI Table 4 and SI Table 5). Together these results highlight the potential for precise control of nanoparticle properties using a biological system.

## Conclusions

Moving forward synthetic biological systems may be a valuable route of nanoparticle production. We demonstrated the ability to program a heterologous host to synthesize arsenic sulfide nanomaterials, and have shown that the characteristics of the resulting materials depend on multiple biological factors. Manipulating the biological factors that impact nanomaterial characteristics using the “control knobs” of synthetic biology has the potential to fine-tune the properties of biogenic nanomaterials, and may even enable the production of materials not possible through conventional methods.

## Experimental procedures

### Bacterial culture conditions

Bacterial strains were transferred from frozen stocks stored at −80 °C and grown overnight under aerobic conditions in 5 ml of lysogeny broth. *Escherichia coli* DH5α strains were grown at 37 °C and *Shewanella* sp. ANA-3 cells were grown at 30 °C in a shaker at 220 rpm. Upon reaching late-log phase in LB, cells were transferred to anaerobic minimal media described here (McFarlane *et al.*, 2015). Concentrations of vitamins, minerals, and amino acids (Table S2) used in the anaerobic minimal media experiments are described in SI. For *E. coli*, 15 mM glucose was added as a carbon source and for *Shewanella* sp. ANA-3 15 mM lactate (ANA-3) was used as a carbon source. 30 mM fumarate was added as terminal electron acceptor for both *E. coli* and ANA-3 cells. Cells grew under anaerobic conditions to reach an OD_600_ of 0.6-0.8.

All the stock solutions (sodium arsenate, sodium metaarsenite, sodium sulfide, sodium thiosulfate) were made anaerobic by flushing the solutions with nitrogen and sterilized by autoclaving them in anaerobic glass serum bottles. Appropriate antibiotics were added to all media to maintain plasmid selection.

### Arsenic sulfide nanomaterials synthesis

For experiments involving microbial reduction of substrates (arsenate and thiosulfate), we induced arsenate reductase genes in *E. coli* (OD_600_ of 0.6-0.8) using 2.5 mM IPTG and 0.5% arabinose, and thiosulfate reductase gene using 2.5 mM IPTG. After 4 hours of induction, we added 5 mM arsenate and/or 2.5 mM thiosulfate to the anaerobic serum bottles.

In experiments to test the influence of cells and supernatant on the size and shape of the arsenic sulfide nanomaterial, late log phase cells of *E. coli* or ANA-3 were centrifuged and washed 3 times with HEPES buffer to remove salts, and cells were then resuspended in buffer and added to sterile serum bottles. For testing cell culture supernatant, cells were filtered using 0.22 µm sterile filter, and then added to sterile serum bottles and oxygen was purged by flushing nitrogen through the system. In case of autoclaved cells or supernatants, cells or cell-free supernatant and oxygen was purged by flushing nitrogen and the bottles were autoclaved. Finally, 5 mM sodium metaarsenite and 10 mM sodium sulfide were added to the anaerobic bottles containing either cells or supernatant. Minimal media controls were tested for abiotic arsenic sulfide synthesis, and arsenic sulfide precipitate was not observed.

All reactions proceeded for 72 hours, and samples were collected and imaged under scanning electron microscopy. Duplicate experiments were performed for each time point. To characterize relative abundance of arsenic sulfide nanomaterial, absorption at 375nm was measured in standard cuvettes using a Nanodrop 2000C (ThermoFisher), as described previously (McFarlane *et al.*, 2015).

### Scanning Electron Microscopy

Arsenic sulfide precipitate was separated from cells and media by centrifugation in 50 ml conical tubes. Precipitates were washed and resuspended, 5 times in DDI water to remove salts. After the washing process, the precipitate was resuspended in a 1 ml DDI water and deposited on a silicon wafer (Ted Pella) and air dried overnight. Samples were coated with gold in a sputter coater (Ted Pella) and scanning electron micrographs were obtained using JEOL 700 FE scanning electron microscopy, accesorized with energy-dispersive X-ray spectroscopy (EDS) for elemental analysis.

### Plasmid construction details

Plasmids expressing the arsenic reduction pathways from *Shewanella sp.* ANA-3 were created using Gibson assembly. *arsDABC* and *arrAB* genes were amplified from *Shewanella* sp. ANA-3 and cloned into plasmids pZE(Reed *et al.*, 2012) and pTSlb(Shis and Bennett, 2013). pZE plasmid was modified for this experiment by replacing kanamycin resistance marker with spectinomycin resistance marker before adding arsDABC gene in place of GFP. arsDABC expression was regulated by a IPTG inducible lac promoter and arrAB was regulated by arabinose inducible pBAD promoter. For nanostructure synthesis, plasmids were transformed into *E. coli* strain JW3470-1 ΔarsC (Baba *et al.*, 2006). This strain has a deletion of the *E. coli* arsenate reductase *arsC*. Note this gene is related but not the same as the *arsC* gene from *Shewanella sp.* ANA-3 we cloned into the pZE arsDABC plasmid. ArsC is an integral protein involved in reduction of arsenate to arsenite; in the absence of this gene, arsenate reduction is inhibited in wildtype *E. coli*. A plasmid containing the thiosulfate reductase gene was obtained from Addgene, pSB74 (https://www.addgene.org/19591/). The gene was under an IPTG inducible promoter and had a carbenicillin resistance marker (Bang *et al.*, 2000b).

### Arsenate reduction analysis

Concentrations of arsenite and arsenate species were measured using ICP-MS (Exova, California). To prepare samples for ICP-MS, *E. coli* cultures were incubated for 72 hours with 5 mM arsenate As(V). Samples were centrifuged and filtered with 0.22 µm filter to remove cells. Arsenate reductase gene, if present, will reduce arsenate As(V) to arsenite As(III). If the cells do not possess arsenate reductase, or the genes were not active, arsenite production would be low or null similar to background levels.

### Thiosulfate reductase activity

*E. coli* strain DH5alpha with pSB74 was tested for thiosulfate reduction. 1 mM thiosulfate was added to cultures after reaching at OD_600_ of 0.7. We measured the activity of thiosulfate reductase gene indirectly, the thiosulfate reductase enzyme converts thiosulfate to sulfide. There is a direct relationship between the gene activity of thiosulfate reductase and sulfide concentration. Cline assay was performed to measure the concentrations of sulfide using the procedure reported in (Bang *et al.*, 2000b). Sulfide concentrations were measured using UV-Cuvette micro cuvettes in a NanoDrop 2000C spectrophotometer at 600 nm, and then concentrations were interpreted by comparing it to the calibrated reference solution of sodium sulfide. Thiosulfate reductase activity was also confirmed by adding 1 mM ferric chloride to the media (Bang *et al.*, 2000a). This is a qualitative assay to confirm the production of sulfide by the bacteria; when expression of the thiosulfate reductase was induced, a black precipitate of iron sulfide was formed. Uninduced cultures formed less precipitate and no precipitate formed in cultures lacking the plasmid containing the thiosulfate reductase gene.

### Nanomaterial size analysis

Nanomaterial dimensions were determined from scanning electron microscopy (SEM) images. Objects in images were measured using ImageJ (Abràmoff *et al.*, 2004). For images with nanofibers, a minimum of 150 fiber widths were measured per image. Three images were measured for each experimental condition.

## Acknowledgements

This work was supported by Office of Naval Research award number N00014-15-1-2573. We thank USC Center for Electron Microscopy and Microanalysis (CEMMA) and Exova for their help with sample anlysis.

